# Multiparent Recombinant Inbred lines crossed to a tester provide novel insights into sources of *cis* and *trans* regulation of gene expression

**DOI:** 10.1101/2025.06.07.658407

**Authors:** Fabio Marroni, Alison M Morse, Adalena V Nanni, Nadja Nolte, Patricka Williams-Simon, Luis Leon-Novelo, Rita M Graze, Paul Schmidt, Elizabeth King, Lauren M McIntyre

**Affiliations:** Department of Agricultural, Food, Environmental and Animal Sciences, University of Udine, Udine, 33100, Italy; Department of Molecular Genetics and Microbiology, University of Florida, Gainesville, FL 32611, USA; University of Florida Genetics Institute, University of Florida, Gainesville, FL 32611, USA; Department of Biotechnology and Systems Biology, National Institute of Biology, Ljubljana 1000, Slovenia; Jožef Stefan International Postgraduate School, Ljubljana 1000, Slovenia; Department of Biology, University of Pennsylvania, 433 S University Ave., 226 Leidy Laboratories, Philadelphia, PA 19104, USA; Department of Biostatistics and Data Science, University of Texas Health Science Center at Houston-School of Public Health, 1200 Pressler St. Suite E809, Houston, TX 77030, USA; Department of Biological Sciences, Auburn University, Auburn, AL 36849, USA; University of Pennsylvania, Department of Biology, 433 S University Avenue, Philadelphia, PA, 19104; Division of Biology, University of Missouri, 401 Tucker Hall, Columbia, MO 65211, USA

## Abstract

We propose crossing multi-parent recombinant-inbred-lines (RILs) to a common tester and measuring allele specific gene expression in the offspring. Testing whether allelic imbalance between two RIL x Tester crosses is equal, is a test of *cis* or *trans* depending on the RIL alleles compared. The study design also enables to separate two sources of *trans* variation, genetic and environmental, detected via interactions with *cis* effects. Examining these components of regulatory variation, we demonstrate this approach in a long-read RNA-seq experiment in female abdominal tissue at two time points in *Drosophila melanogaster*. Among the 40% of all loci that show evidence of genetic variation in *cis, trans* effects due to the environment are detectable in 31% of loci and *trans* effects due to genetic background are detectable in 19% of loci with little overlap in sources of *trans* variation. The loci identified in this study are associated with loci previously reported to exhibit genetic variation in gene expression in a range of tissues and large population samples, suggesting that there is consistent variation for genetic regulation of gene expression. We show that eleven loci in a QTL for thermotolerance, previously shown to differ in expression based on temperature, have evidence for regulation of gene expression regardless of the environment, including Cpr67B, a cuticular protein suggesting a potential functional role for standing variation in gene expression. This study provides a blueprint for efficiently identifying regulatory variation in gene expression, as the tester design maximizes *cis* variation and enables the efficient assessment of all pairs of RIL alleles relative to the tester, a much smaller study compared to the pairwise direct assessment.

## Introduction

Spatial and temporal control of gene expression is known to be ‘remarkably complex’ (Arnone and Davidson 1997). The evolution of these complex components of gene expression regulation has been a subject of lively discussion since we have been able to empirically examine these questions (McGinnis et al. 1984; Scott and Weiner 1984; Krumlauf 1994). This involves unpacking the relative contributions of evolutionary changes in different components, for example contrasting protein coding versus *cis*-regulatory regions (Stern and Orgogozo 2008; Durkin et al. 2024) or *trans* regulatory factors; *cis* regulatory variation in promoters and enhancers, or *cis* by *trans* interactions (Mattioli et al. 2020; Ballinger et al. 2023; Hansen et al. 2024). Allelic imbalance as a method to empirically estimate genetic variation in *cis* and *trans* regulation was laid out in a seminal paper in 2004 (Wittkopp et al. 2004). Tests for whether the source of differences in allelic expression are due to *cis* effects are usually tests between alleles within an individual, while to detect *trans* effects an allele’s expression is typically compared between individuals (e.g. Wittkopp et al. 2004; Fear et al. 2016; Osada et al. 2017). *Trans* tests between individuals have lower power than the within individual *cis* effect tests for allelic imbalance and power can be further reduced if differences in overall expression between the environments is ignored (León-Novelo et al. 2014). The power of the test of allelic imbalance depends upon the magnitude of the difference between alleles, the number of replicates and the level of expression (Sherbina et al. 2021). Differences in allelic imbalance in genetically identical individuals compared between environments indicates that *trans* variation due to the environment is differentially interacting with the *cis* polymorphisms; presence of the *cis* polymorphisms is a prerequisite to discriminate the two alleles being expressed (Fear et. al. 2016, León-Novelo et al. 2018).

Mapping eQTL for overall expression identifies ‘local’ and ‘distal’ sources of variation in overall gene expression (Rockman and Kruglyak 2006). eQTL studies test that the source of the expression variation is genetic, as in QTL studies. All QTL studies require hundreds of samples to map genetic variation. For eQTL there is an obvious link between expression at a locus and the regulatory variation at that locus (*cis*) but the typical test averages many lines to contrast alleles and will include all of the *trans* effects and the interactions (Figure 1). The test of mean overall expression differences between a pair of inbred lines (*e*.*g*. between the two parental lines) is the same as the test of allele imbalance when there is an absence of *trans* regulatory effects and their interactions contributing to expression variation (Fear *et. al*. 2014). Direct comparisons between individual RIL lines can also be used to infer *cis* and *trans* effects and these between genotype comparisons also involve interactions (Genissel et al. 2008).

**Figure 1:**
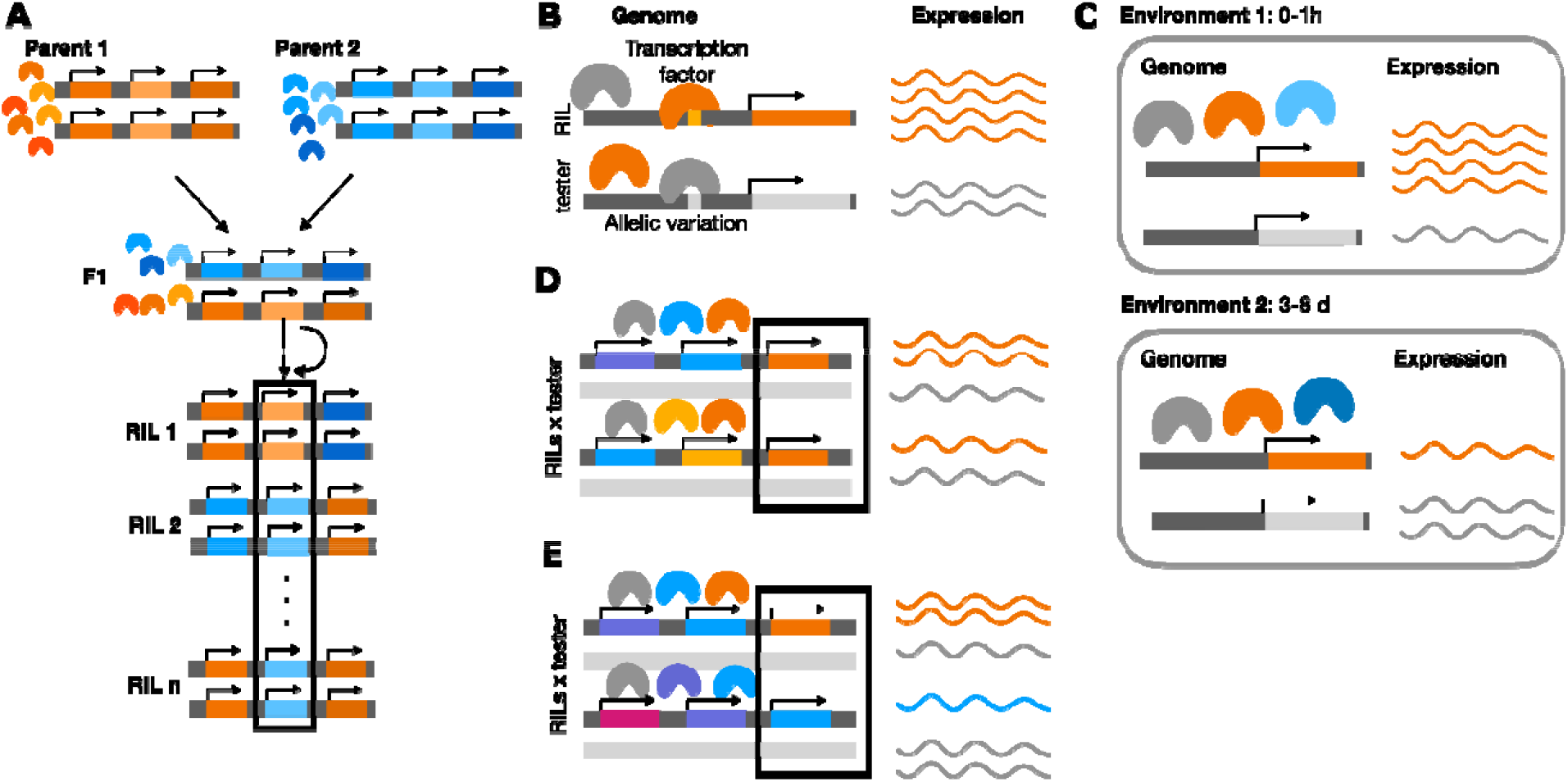
*Cis-* and *trans-* sources of regulatory variation in tests of allelic imbalance. **A:** Comparing expression of the two alleles between two parents (*cis, trans* and interactions), within an individual F1 (*cis* and interactions but no main effect of *trans* since the environment is shared) or as an average between RILs (local, distal and interactions). **B:** Comparing allelic imbalance within an individual in a testcross or (: *cis*) **C:** between different conditions for the same genotypes (); differences result from environmental changes in *trans* regulation (: *trans* environment). **D:** Comparing RIL x tester genotypes. When the RIL allele is the same between two crosses, differences are due to *trans* (: *trans* genetic). **E:** Comparing RIL x tester genotypes. When the RIL allele is not shared, (: *cis* genetic). All tests include effects of *cis by trans* interactions.

We propose a genetic design with crosses between multi-parent recombinant inbred lines (RILs) and an inbred tester with a different genetic background from the RIL parents. This cross produces genotypes that are heterozygous at most loci. When there is allelic imbalance in expression within the individual there are *cis* regulatory polymorphisms, and the potential for variation in *trans* to interact with the regulatory polymorphisms in *cis*. The test of the null hypothesis that the allelic imbalance *between* two RIL x Tester crosses is equal compares individuals to determine if the imbalance differs between them. Given the presence of *cis* variation, magnified by the choice of the tester, additional *cis* or *trans* sources of regulatory variation can be identified. Here, unlike in the other scenarios, the test statistic is the same for the effects of *cis* and the effects of *trans* as the interpretation of the test depends on the alleles of the RILs at the locus (Figure 1). To our knowledge, this is a novel proposal that enables testing for *cis* and *trans* effects for the same genes and alleles, using the same framework and without confounding the choice of the test statistic with the resulting inferences.

A multiparent RIL crossed to a tester, leverages extensive investments in multi-parent genetic resources. Multi-parent genetic resource panels have been developed for many plant breeding applications in many different systems including maize (Dell’Acqua et al. 2015), rice (Raghavan et al. 2017), wheat (Mackay et al. 2014), sorghum (Ongom and Ejeta 2018), barley (Sannemann et al. 2015), soybean (Hashemi et al. 2022), strawberry (Mangandi et al. 2017), and okra (Sandeep et al. 2025). Multi-parent populations have also been developed for QTL mapping in many model organisms such as Arabidopsis (Kover et al. 2009), Drosophila (King et al. 2012), *C. elegans* (Noble et al. 2017), mouse (Srivastava et al. 2017), and yeast (Cubillos et al. 2013).

Since the first results in QTL mapping (Lander and Botstein 1989), the field has grappled with how to move from QTL regions to direct links between genes and phenotypes. There have also been proposals to combine QTL/GWAS, and eQTL to uncover regulatory networks underpinning complex traits (e.g. Jansen et al. 2009; Zhu et al. 2016). Here we explore the idea of moving from QTL to candidate genes by overlaying information about standing variation in regulation of gene expression. Using the results of a QTL study on thermotolerance, we selected two multi-parent RIL lines from each of the alleles in the main QTL for thermotolerance for this study (Williams-Simon et al. 2024). We conducted a replicated long read RNA-seq experiment in *Drosophila melanogaster*, where we measure allelic imbalance. We crossed four multi-parent RIL lines to a common tester, and by comparing pairs of RIL x Tester crosses we identify genetic sources of regulatory variation in *cis* and *trans*. A comparison between two time points for abdominal tissue identifies environmental sources of *trans* effects. We integrate the results of the present work with previous QTL mapping (Williams-Simon et al. 2024).

We used a Bayesian model that accounts for difference in overall expression, as well as potential differences in map bias between alleles (León-Novelo et al. 2018). We show how the initial inferences about expression differences between the parental alleles are both supported and enriched by this study(Williams-Simon et al. 2024). Overall, we find that RIL lines crossed to a tester provide additional novel insights into the relative sources of *cis* and *trans* regulatory variation.

## Methods

For a QTL analysis of thermotolerance, the most significant Quantitative Trait Loci (QTL) region identified by Williams-Simon et al. (2024) was used to randomly select two RILs with the shared allele for ‘high’ thermotolerance in QTL 1 (A5: dm12272, dm12279) and two RILs with the ‘low’ thermotolerance allele (A4 : dm11037, dm11255).

### Fly Husbandry

Abdominal tissue was obtained as follows: populations were housed in narrow vials (Genesee Scientific – 32-116SB) at 25°C and 60% relative humidity, on a modified Bloomington Cornmeal Recipe (Beltz et al. 2025). The four RILs selected from the *Drosophila* Synthetic Population Resource (DSRP) (King et al. 2012) were crossed with a common female genotype W1118 (Bloomington *drosophila stock center stock number 3605*). Each cross consisted of 10 female W1118 and 5 RIL males, individuals were placed in a common vial as pupae. 72 hours after mating 60-70 F1 eggs were transferred to a new vial with fresh food. At 10 days post oviposition, 0-1 hour virgin females were collected by observing eclosion in 1 hour increments. Newly eclosed females were either dissected for abdominal tissue and placed in RNAlater (Sigma – MKCJ9161) + 100uL 0.1% PBST and at -20°C, or transferred to a new vial with fresh food. After 3-8 days, ovaries were removed (gonadectomized) and flies placed in RNAlater and stored at -20°C. Three independent replicates of ∼50 individuals were collected for each cross and time point.

### Library Preparation and Sequencing

For each sample, mRNA was isolated (DynaBeads mRNA direct kit) from a pool of ∼20 abdomens. ONT libraries were constructed using the ONT PCR-cDNA Barcoding Kit (SQK-PCB109) starting with polyA mRNA according to the manufacturer’s protocol. Libraries were pooled to a total of 100 fmol and run on a MinION Mk1c with real-time basecalling and demultiplexing (Guppy v6.1.5, MinKNOW v22.05.8). Read length and quality was evaluated with all samples passing the PycoQC metrics v2.5.2 (Leger and Leonardi 2019). Based on the MinION read counts, libraries were repooled prior to obtaining additional sequencing data on the ONT PromethION (Guppy v5.1.13, MinKNOW v23.04.5) at the University of Florida Interdisciplinary Center for Biotechnology Research (ICBR). Technical replicates (TRs) are defined as the same library run on different ONT flow cells (MinION or PromethION). TRs 1-3 were run on the MinION and TRs 4-6 were run on the PromethION.

Fast5 files were converted to pod5 formats (pod5 v 0.3.6), and basecalling was executed using Dorado (v 0.5.2) (https://github.com/nanoporetech/dorado) with options --recursive --device “cuda:0,1” --kit-name SQK-PCB109 --trim none. Reads were demultiplexed using the demux mode of Dorado (v 0.5.2) with options --no-classify –emit-fastq. Fastq files are available at the SRA under BioProject PRJNA1134728. The fastq files generated by were processed using pychopper (v 2.7.1), the oriented fastq files were aligned to *D. melanogaster* 6.50 and the resulting sam files were converted to gtf using samtools (v 1.10) (Li et al. 2009) and bedtools (v 2.29.2) (Quinlan and Hall 2010). The resulting gtf files (67 technical replicates from 24 samples), the *D. melanogaster* 6.50 fasta reference file (https://ftp.flybase.net/releases/FB2023_01/dmel_r6.50/ (Öztürk-Çolak et al. 2024)), and a design file were used as input to SQANTI-reads in order to calculate metrics used for quality control (Keil et al. 2025).

### Construction of haplotype specific references

SNP data for w1118 and the RILS were retrieved from the *Drosophila* Synthetic Population Resource data (King et al. 2012; Fear et al. 2016); genomic coordinates were converted from release 5 (Hoskins et al. 2007) to release 6 (Hoskins et al. 2015) using the coordinate converter tool in Flybase (Gramates et al. 2022). As part of quality control, 11, 5, 6, and 6 SNP positions were discarded in dm11037, dm11255, dm12272, and dm12279 respectively, because the reference base in the VCF file did not match the reference base in the genome.

The dmel6 reference genome (Hoskins et al. 2015) was updated with w1118 and RIL specific SNPs respectively (Graze et al. 2012; Munger et al. 2014; Gobet et al. 2022). Initial alignments showed some evidence of systematic bias toward the RIL genotypes. We evaluated the alignments at positions expected to be heterozygous for a RIL SNP. If there were a min of 5 reads and no evidence of the expected polymorphism among the reads, the SNP was presumed to be a missed call in w1118 and the w1118 VCF file was updated to include the RIL SNP (available at: https://github.com/McIntyre-Lab/papers/blob/master/marroni_2025/VCFs/). The amended w118 and the RIL vcf files were used to update the reference genome. Haplotype specific gene regions (+/-100 base pairs from the annotated transcription start and end site) were identified. Reads were mapped separately to the parental haplotype specific gene regions using minimap2 (Li 2018) with the following parameters: -a -x splice --secondary=yes -N 200 -p 0.9 -C 5. Uniquely mapping reads were compared and identified as allele specific if they mapped with fewer mismatches to one parent (Graze et al. 2012) using the python script sam_compare_w_feature.py (Miller et al. 2021) available at https://github.com/McIntyre-Lab/BayesASE) and no haplotype bias was observed. The resulting allele counts together with the non-allele specific gene counts are used as input into the Bayesian Model. Genes with a prior of 0, or with fewer than 10 allele specific reads were excluded from analysis.

### Statistical Tests

To detect differences in allelic imbalance we used the Bayesian analysis of allele-specific expression (BayesASE) based on the *environmental model* (León-Novelo et al. 2018; Miller et al. 2021). In environment/condition 1 the proportion of reads produced by the reference allele is *θ*_1_ and allelic imbalance in the genotype/condition 1 occurs when *θ*_1_ ≠ 1/2. *θ*_2_ is defined analogously to *θ*_1_ but in environment/condition 2. There are three hypothesis tests based on the model: allelic imbalance in condition 1 (*H*_01_: *θ*_1_ = 1/2 vs *H*_01_: *θ*_1_ ≠ 1/2); allelic imbalance in condition 2 (*H*_02_: *θ*_2_ = 1/2 vs *H*_02_: *θ*_2_ ≠ 1/2); allelic imbalance differs across the two conditions (*H*_03_: *θ*_3_ =0 vs *H*_03_: *θ*_3_ ≠0 with *θ*_3_ = *θ*_1_ - *θ*_2_). A pre-condition for testing *H*_03_ is the significant effect of allelic imbalance in at least one cross (rejection for *H*_01_ or H_02_).

We briefly summarize the *environmental model* of León-Novelo et al (2018). Inference here is at the gene level, that is the model is fitted separately for each gene. For each gene, we have the set of counts {(*x*_*ik*_, *y*_*ik*_, *z*_*ik*_):*i* = 1,2; *k* = 1,…, *K*_*i*_ } mapping to the RIL (*x* counts), tester (*y*) and both alleles (*z*). Here *i*= 1,2 indexes the condition (in our context the two RIL x tester crosses being compared, or, in the second experiment, the same RIL x tester at a different developmental time) and *k*= 1, …, *K*_*i*_ indexing the number of independent replicates with *K*_*i*_ in condition *i* (in our experiments *K*_*i*_ = 3). The model assumes that *r*_*i,RIL*_ n and *r*_*i,tester*_ are known, where the former is the probability that a read generated by the RIL aligns to the RIL (as opposed to aligning to both) and the latter is defined analogously replacing the RIL by the tester; when *r*_*i,RIL*_ ≠ *r*_*i,tester*_ there is mapping bias. These probabilities can differ across conditions *i* = 1,2 and across genes. We estimate these with the Prior Calculation module from BayesASE. We assume the counts are generated from a negative binomial distribution. More specifically, *x*_*ik*_ ∼ *NB(α*_*i*_ *β*_*ik*_ *r*_*i,RIL*_, *φ*), *y*_*ik*_ ∼ *NB*((1/*α*_*i*_) *β*_*ik*_ *r*_*i,tester*_, *φ*) and 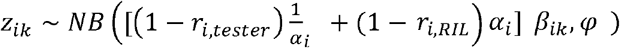 where *NB*(*µ, φ*) denotes the negative binomial distribution with mean *µ* and dispersion parameter φ, thus with variance µ+ *φµ*^2^. The replicate specific random effect *β*_*ik*_ accounts for differences in coverage between replicates (A replicate with higher coverage will tend to have larger (*x, y, z*) counts than one with lower coverage); *r*_*i,RIL*_ and *r*_*i,tester*_ accounts for mapping bias and *α*_*i*_ ∈ (0, ∞) is the parameter related with the level of allele-imbalance in condition *i*, with *α*_*i*_ =1 indicating no imbalance. The proportion of reads coming from the RIL allele under condition *i* is

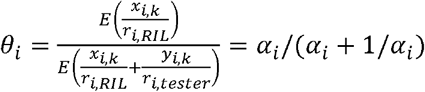

Inference is based on the central, in our case 95%, CrIs of *θ*_1_, *θ*_2_ and *θ*_3_ = *θ*_1_ - *θ*_2_. The priors for the model parameters: 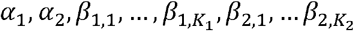 and *φ*, and more detailed interpretation of the parameters are given in detail in León-Novelo et al (2018). For each gene, we estimated the probability of a read aligning uniquely to the allele that generated it from the RNA long-read data using the Prior Calculation module from BayesASE (Miller et al. 2021). All comparisons were carried out with 100,000 iterations and a burn in of 10,000. The null hypotheses *H*_01_, *H*_02_ are rejected if the CrIs for *θ*_1_, *θ*_2_ do not contain 0, respectively. *H*_03_ is rejected if either *H*_01_ or *H*_02_ is rejected *and* if the CrI for *θ*_3_ does not contain 0. The tests of *θ*_1_/*θ*_2_ are within individuals and the test of *θ*_3_ is a comparison between individuals. We classify our inferences for *θ*_1_, *θ*_2_ as *cis* effects (H_01_ *cis*), because in tests within individual there are no differences in *trans*. The interpretation of the effects of *θ*_3_ depends on the context (Figure 1). Results of the Bayesian model used for testing are available on github (https://github.com/McIntyre-Lab/papers/blob/master/marroni_2025/bayesian_out).

### Classification of *cis* and *trans* results from H_03_

There are three comparisons we make using *θ*_3_. All three comparisons use the same statistical approach, removing confounding of inferences about the magnitude of effects with the use of different test statistics. We label these three comparisons: *cis* genetic between individuals, *trans* genetic between individuals, and *trans* due to the environment. However, we note that because the test of *θ*_3_ depends upon *θ*_1_/*θ*_2_ a rejection of the null by definition includes an interaction between that effect and the *cis* differences between the RIL and tester alleles.

When the alleles for the two RIL x tester crosses compared differ for gene of interest, *H*_03_ is a test of *cis* and labeled as *H*_03_ *cis*. When the alleles for the two RIL x tester crosses compared are shared, but the RIL background differs, the *cis* genotype is the same in both conditions and H_03_ is a test of genetic differences in the background of the two RIL lines, labeled as *H*_03_ *trans* genetics effect. When the same RIL x tester cross is compared at the two developmental time points, *H*_03_ is a test of environmental difference labeled as *H*_03_ *trans* environment effect (Figure 1).

### Comparison with published evidence of genetic variation

Genes with evidence for allele imbalance in this study were compared with genes showing heritability of gene expression (Wayne et al. 2007); *cis* and *trans* effects in comparisons among RI lines of OregonR and 2b3 (Genissel et al. 2008), *cis* effects as estimated by allelic imbalance within an individual in a testcross with W1118 and DGRP lines (León-Novelo et al. 2018) and *cis* effects identified in F1’s of interspecific hybrid crosses (Graze et al. 2012). Comparisons were performed as a Fisher’s exact test of independence between the binary indicators of significance from this study and the published literature (Rivals et al. 2007).

## Results and Discussion

In order to test for allelic imbalance, polymorphisms in the exonic regions of the gene need to be captured enabling the assignment of reads to alleles. Early studies of allele imbalance relied on large differences between the alleles for this reason. In this study, almost all of the genes in *D. melanogaster* 6.50 (Öztürk-Çolak et al. 2024) had at least 1 SNP in the exonic regions for either the tester (W1118) or the RIL. There were only 538, 614, 498, 548 genes (out of 16,834) in dm12279, dm12272, dm11255, dm11037 respectively that could not be evaluated for allelic imbalance due to lack of polymorphisms between the tester (W1118) and RIL haplotype.

### Long read sequencing

We generated 128,001,025 million long reads. The median number of reads per sample was 4,671,327 with ∼11,000 genes detected (at least one read) and ∼5,700 genes with at least 25 reads (**Table S1**). The dm12279 replicate 1 sample at 0 to 1 hour had a lower proportion of mapped reads compared to the other samples, all other samples were found to be of similar and good quality (**Table S1)**. A median of 3910 genes satisfied the conditions for being tested for allelic imbalance (**Table S2**). ***cis* and *trans* sources of regulatory variation**. The proportion of genes showing allelic imbalance in the test *H*_01_ (within individual cross) is 28% and across all 4 RIL x Tester crosses in both environments we detect *cis* effects in ∼39% of genes (**Table S4**). The unique property of this design is that every test of *H*_03_ between a pair of RIL x Tester crosses is the same statistical test, but the inference about the source of regulatory variation differs. We condition the tests between individuals (*H*_03_) on the test of *H*_01_ between the RIL and the tester. When the RIL alleles differ between crosses, the test between crosses is a test of *cis* differences between the RIL alleles. This test will also include *trans* effects from the different RIL backgrounds that interact with *cis* differences among the three alleles. We see evidence for *cis* differences between RIL alleles at 63% of the genes tested. When the RIL haplotype is shared between individuals, the *cis* effects are shared between the two crosses, as in a reciprocal cross. Differences in allelic imbalance between RIL x Tester crosses with the same genotype at the expressed locus are due to *trans* sources from the genetic background, and this test will also include interactions with *cis* differences among the three alleles. We find evidence for a genetic effect in *trans* for 19% of the genes tested. The test statistic used in the test of *cis* and *trans*, is the same and differences are unlikely to be the result of the test itself. Interestingly, all but 57 of the genes with evidence for *trans* also have evidence for *cis*, suggesting that allelic interactions between *cis* and *trans* sources of variation are prevalent **(Figure 2**). In addition, only 48 genes showing *trans* variation due to environment show *trans* variation due to genotype (**Figure 2**).

**Figure 2:**
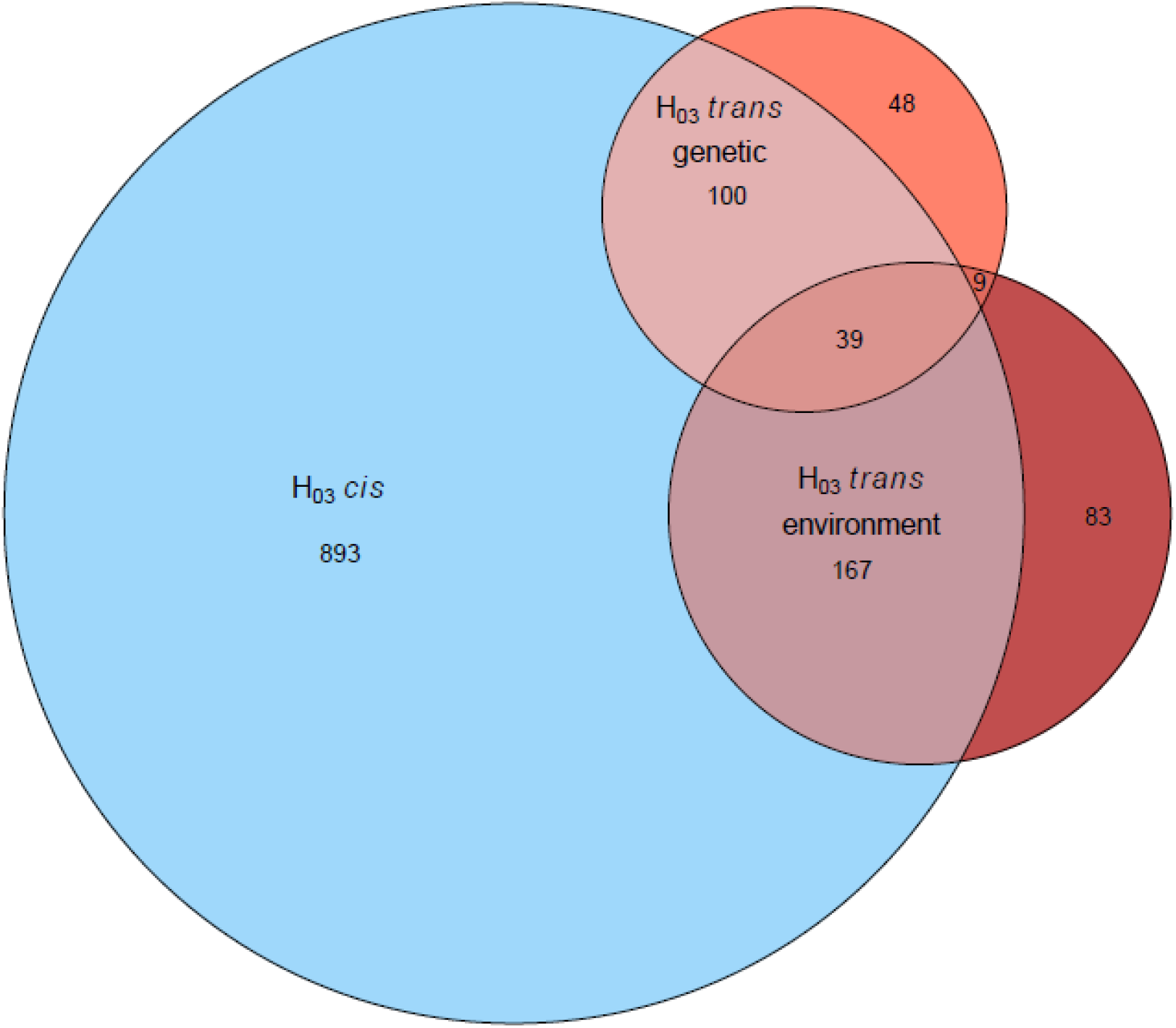
Venn diagram showing the overlap of genes for which was rejected when comparing different genotypes sharing the RIL allele being tested (*trans* genetic), the same genotype in different times (*trans* environment) and different genotypes not sharing the RIL allele being tested (*cis* genotypic).

We examined sources of *cis* and *trans* for each pair of RIL x Tester crosses. We see a consistent pattern (more *cis* than *trans*) for each genotype and in both environments (**Figure 3**). We also see consistent estimates of the relative proportion of *cis* and *trans* for both the higher coverage in 3-8 days and lower coverage 0-1 hours, although consistent with expectations higher coverage improves detection of both sources of regulatory variation.

**Figure 3:**
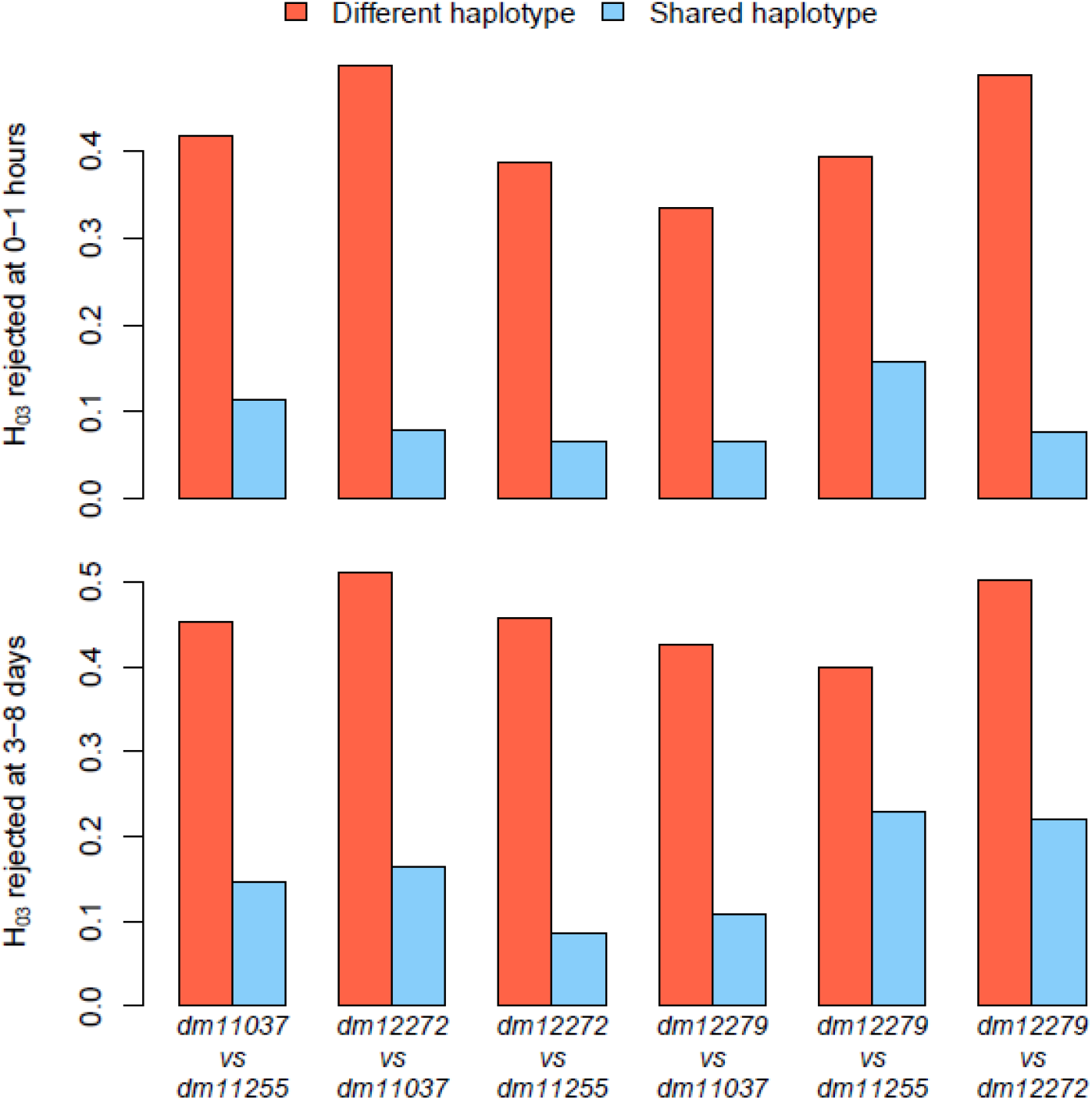
Proportion of genes showing different levels of allelic imbalance between lines (Rejection of), limited to regions in which the lines have different (red) or same (blue) haplotype.

### The environment is a *trans* effect

The difference in presence/absence of gene expression is also expected to be due to *trans* factors attributable to external conditions or, like in our study, developmental stage. We generically term this contribution of environmental conditions as *environment*. There were 16,834 genes in the annotation. Of these 13,804 were detected in either environment (0-1 hours) or (3-8 days). In order to account for the difference in coverage between the 0-1 hour and the 3-8 days samples, we used the genes detected in the 0-1 hour that are absent in the 3-8 days samples as an estimate of *trans* presence/absence (*n=355*). Assuming the effect is symmetric between the two time points, this would suggest that ∼700 genes or 5% of genes differ in presence/absence *trans* effects. This relatively small difference in presence/absence compared to many tissue comparisons, is likely due to the similarity of the two environments. For loci expressed at both time points with evidence of allele imbalance at either time point, we estimate that (320) 31% of the loci show *trans* effects due to the environment. There is little overlap between loci with *trans* genetic effects and *trans* environment effects. Of the 561 loci that could be evaluated for both effects, 9% showed evidence for both, and 42% showed evidence for either. Yet, it is noteworthy that even though the environment here is similar and these effects are likely a lower bound on the *trans* environmental variation, the effect is much larger than the *trans* genetic effect.

### Candidate loci for thermotolerance

Using gene expression of head tissue, Williams-Simon and colleagues identified genes located in a major QTL for thermotolerance, that differed in expression between high and low RILs in head tissue (Williams-Simon et al. 2024). These genes were considered functional candidates for thermotolerance. The use of ASE results (although in a different tissue) provides further evidence of the existence of genetic effects on regulatory variation of these genes, strengthening their potential role as candidates in thermotolerance. If one of these genes showed evidence for genetic regulatory variation (*cis* or *trans*) and an absence of *trans* effects between the environments, we consider this additional evidence of genetic regulatory variation worth considering in potential follow-up studies for understanding thermotolerance. There are 11 genes that meet these criteria (**Table S3**) including FBgn0035985, Cpr67B. This gene makes a cuticular protein, and is expressed in both the head and the crop, hindgut and rectal pad according to FlyAtlas (Krause et al. 2022). The gene contains an Rebers and Riddiford (R&R) motif (Rebers and Riddiford 1988), which can bind chitin in some circumstances (Togawa et al. 2004). FBgn0035985 further shows significantly different levels of ASE between low and high lines, further reinforcing its credibility as a candidate for thermotolerance. Another gene, FBgn0039719 (CG15515) is predicted to be a structural component of chitin-based larval cuticle in FlyBase version FB2025_03 (www.flybase.org). FBgn0004387, Klp98A is a kinesine-like protein, which has been shown to be involved in asymmetric division (Derivery et al. 2015). The function is not apparently related to thermotolerance. An additional study integrating GWAS and transcriptome analysis to investigate thermotolerance in *D. melanogaster* (Lecheta et al. 2020) identified the gene as being differentially expressed in a comparison between heat shocked and control *D. melanogaster*, although the gene did not present evidence of association in the GWA analysis. FBgn0035983, CG4080 is suggested to have MHC binding activity, zinc ion binding activity and ubiquitin binding activity, all functions that do not seem strictly linked to thermal tolerance. However, the gene has also shown differential expression as a consequence of both heat and cold shock (Lecheta et al. 2020), and has been associated to thermal sensitivity in *D. melanogaster* (Soto et al. 2025), thus representing an interesting candidate. FBgn0015577, alpha-Est9 was differentially expressed as a consequence of a heat shock (Lecheta et al. 2020). FBgn0039656, CG11951, and for FBgn0266579, tau, showed differential expression as a consequence of heat and cold shock (Lecheta et al. 2020). How the remaining four genes identified (FBgn0000533, FBgn0011769, FBgn0039543, FBgn0086346) may be involved in thermotolerance is less obvious.

## Conclusion

Studies of allelic imbalance test for the presence of regulatory variation in gene expression between alleles at a particular locus. Further focusing on loci with regulatory effects on expression that are not influenced by the environment increases the chances of identifying stable genetic regulatory effects. However, regulation of gene expression is only one of many possible sources of phenotypic variation for any complex trait. Hunting for individual candidate genes and testing them for phenotypic effects assumes that there are large effects loci underneath the QTL worth identifying. There are many cases for which this has been a successful strategy. However, the importance of large numbers of small effect polygenes to quantitative variation remains a potent alternative hypothesis. This hypothesis has been discussed in many contexts since its initial proposal (Jacob 1977). One factor contributing to the ongoing difficulty in evaluating these hypotheses is the lack of efficient designs that enable a survey of a large number of alleles and provide a statistically powerful approach to interrogating *cis* and *trans* effects. The RILxTester design we propose here enables the evaluation of **n** alleles in **n** crosses and by comparing these crosses pairwise can be used to compare the **n** alleles for both *cis* and *trans* effects.

We compared our results to other studies of genetic variation in gene expression (**Table S5**). The genes associated with *cis* effects in this study, were enriched for genes showing evidence for heritability of gene expression in a diallel (Wayne et al. 2007); *cis* and *trans* effects in comparisons among RI lines of OregonR and 2b3 (Genissel et al. 2008), *cis* effects as estimated by allelic imbalance within an individual in a testcross with W1118 and DGRP lines (León-Novelo et al. 2018) and *cis* effects identified in F1’s of interspecific hybrid crosses (Graze et al. 2012). This is a remarkable consistency of evidence for genetic variation in regulation of gene expression, across platforms (microarray, short read RNA-seq and long read RNA-seq), tissue types (head, whole body, abdomen), and experimental designs. This could be because different variants at the same locus contribute in multiple studies due to high levels of standing variation. Relatively stable variation at the same loci could be explained by balancing selection, relaxed selection or variation maintained by mutation-selection balance under stabilizing selection. Another possibility is that the same variants are contributing, with allelic variants that impact expression producing a signal that in combination with the statistical approach is unusually robust to tissue, environment or other microenvironmental variation. In either case, this is support for the hypothesis that genetic regulation of gene expression may be one mechanism by which pleiotropic effects are regulated.

## Data Availability

Fastq files are deposited to the SRA BioProject PRJNA1134728. Results from the Bayesian model are available in individual files labeled by the test performed, and can be accessed at https://github.com/McIntyre-Lab/papers/blob/master/marroni_2025/bayesian_out. The individual vcf files used for updating are available at https://github.com/McIntyre-Lab/papers/blob/master/marroni_2025/VCFs. We are also attempting to deposit the pod5 files (raw oxford nanopore sequencing data) but they are too large for zenodo and we are struggling to find a repository.

## Funding

This work was funded by GM GM137430 (LM, PS), R01GM128193 (LM) R35 GM149238 (EK), R35 GM133376 (RR), the Fund for the National Research Program and Projects of Significant National Interest (PRIN), CUP: G53D23002620006, 2022E8NN2N (FM), the National Science Foundation Career award DEB1751296 (RG), and the LongTrec European Marie Curie Action Network (NN).

## Acknowledgements

The Department of Molecular Genetics and Microbiology, The University of Florida Cancer Center, The University of Florida Genetics Institute, University of Florida Research Computing, HiPerGator.

## Notes

### Competing Interest Statement

The authors have declared no competing interest.

### Summary of Updates

The manuscript was changed from a brief version to a full length investigation. As such, a more detailed methods section has been included. In addition, Figures were improved, to make them clearer.

