## Supplementary File 1 for "Multiparent Recombinant Inbred lines crossed to a tester provide novel insights into sources of *cis* and *trans* regulation of gene expression": README.docx

These files are the output of the Bayesian model for detection of allelic imbalance.

bayesian_out_dmYYYYY_XXZ.csv: output file measuring AI in one condition

bayesian_out_dmYYYYY_01h_vs_dmZZZZZ_38d.csv: output file measuring AI between different timepoints in the same line

bayesian_out_dmYYYYY_XXX_vs_dmZZZZZ_WWW_01h.csv: output file measuring AI between different lines at the same timepoint

Description of field for files of the type bayesian_out_dmYYYYY_XXZ.csv

| **Column name** | **Column meaning** |
| --- | --- |
| Comparison | name of the comparison |
| FEATURE_ID | name of the gene |
| dm11037_01h_num_reps | number of replicates |
| counts_dm11037_01h_g1 | Number of reads assigned to haplotype 1 (g1) |
| counts_dm11037_01h_g2 | Number of reads assigned to haplotype 2 (g2) |
| counts_dm11037_01h_both | Number of reads not assigned to haplotypes |
| prior_dm11037_01h_g1 | Prior probability of a read mapping to haplotype 1 |
| prior_dm11037_01h_g2 | Prior probability of a read mapping to haplotype 2 |
| H3_independence_Bayes_evidence | pvalue for H3 violation (not used for one condition) |
| dm11037_01h_sampleprop | Proportion of reads assigned to haplotype 1: g1/(g1+g2) |
| dm11037_01h_theta | Estimated proportion of reads assigned to haplotype 1 |
| dm11037_01h_q025 | LCL for proportion of reads assigned to haplotype 1 |
| dm11037_01h_q975 | UCL for proportion of reads assigned to haplotype 1 |
| dm11037_01h_Bayes_evidence | pvalue for AI |
| dm11037_01h_AI_decision | Was AI detected? 1 = Yes, 0= No |
| alpha1_postmean | Posterior mean of alpha |
| flaganalyze | Should this gene be analyzed? 1 = Yes, 0= No |

Description of field for files of the type bayesian_out_sample_1_vs_ sample_2.csv

| **Column name** | **Column meaning** |
| --- | --- |
| comparison | name of the comparison |
| FEATURE_ID | name of the gene |
| sample_1_num_reps | number of replicates in sample1 |
| sample_2_num_reps | number of replicates in sample2 |
| counts_sample_1_g1 | Number of reads assigned to haplotype 1 (g1) in sample 1 |
| counts_sample_1_g2 | Number of reads assigned to haplotype 2 (g2) in sample 1 |
| counts_sample_1_both | Number of reads not assigned to haplotypes in sample 1 |
| counts_sample_2_g1 | Number of reads assigned to haplotype 1 (g1) in sample 2 |
| counts_sample_2_g2 | Number of reads assigned to haplotype 2 (g2) in sample 2 |
| counts_sample_2_both | Number of reads not assigned to haplotypes in sample 2 |
| prior_sample_1_g1 | Prior probability of a read mapping to haplotype 1 in sample 1 |
| prior_sample_1_g2 | Prior probability of a read mapping to haplotype 2 in sample 1 |
| prior_sample_2_g1 | Prior probability of a read mapping to haplotype 1 in sample 2 |
| prior_sample_2_g2 | Prior probability of a read mapping to haplotype 2 in sample 2 |
| H3_independence_Bayes_evidence | pvalue for H_03_ violation (if <0.05 H_03_ is violated) |
| sample_1_sampleprop | Proportion of reads assigned to haplotype 1 in sample 1: g1/(g1+g2) |
| sample_1_theta | Estimated proportion of reads assigned to haplotype 1 in sample 1 |
| sample_1_q025 | LCL for proportion of reads assigned to haplotype 1 in sample 1 |
| sample_1_q975 | UCL for proportion of reads assigned to haplotype 1 in sample 1 |
| sample_1_Bayes_evidence | pvalue for AI in sample 1 |
| sample_1_AI_decision | Was AI detected in sample 1? 1 = Yes, 0= No |
| sample_2_sampleprop | Proportion of reads assigned to haplotype 1 in sample 2: g1/(g1+g2) |
| sample_2_theta | Estimated proportion of reads assigned to haplotype 1 in sample 2 |
| sample_2_q025 | LCL for proportion of reads assigned to haplotype 1 in sample 2 |
| sample_2_q975 | UCL for proportion of reads assigned to haplotype 1 in sample 2 |
| sample_2_Bayes_evidence | pvalue for AI in sample 2 |
| sample_2_AI_decision | Was AI detected in sample 2? 1 = Yes, 0= No |
| alpha1_postmean | Posterior mean of alpha in sample 1 |
| alpha2_postmean | Posterior mean of alpha in sample 2 |
| flaganalyze | Should this gene be analyzed? 1 = Yes, 0= No |
